## Supplemental Figures for "Processive and stochastic CHH methylation by plant DRM2 and CMT2 revealed by single-read methylome analysis"

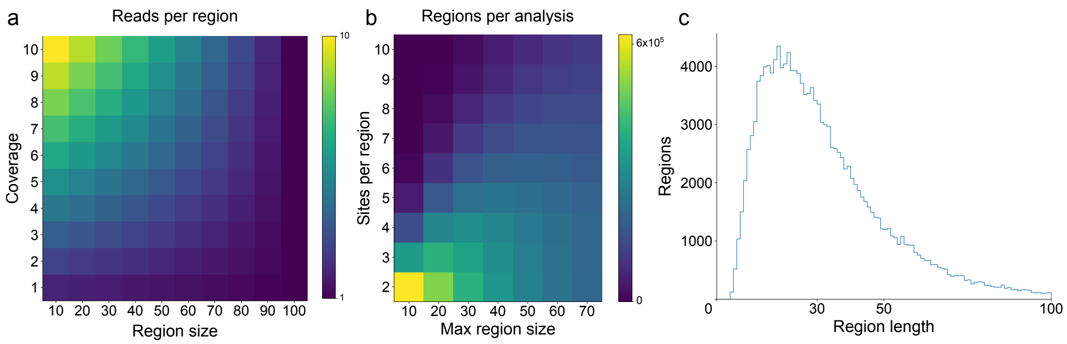


Figure S1: **Technical and biological limitations of region selection.** (a) Expected average number of reads overlapping regions based on region size and library coverage (per strand), given a read length of 100bp. (b) Number of regions in analyses of a wild type *A. thaliana* sample; analysis parameters (max region span in base pairs and exact number of CHH sites) are indicated on the axes. Regions shown have a minimal level of methylation (>= 5%). (c) Distribution of region span of an analysis of the same sample as in (b), with 5 CHH sites and a maximal span of 100bp.

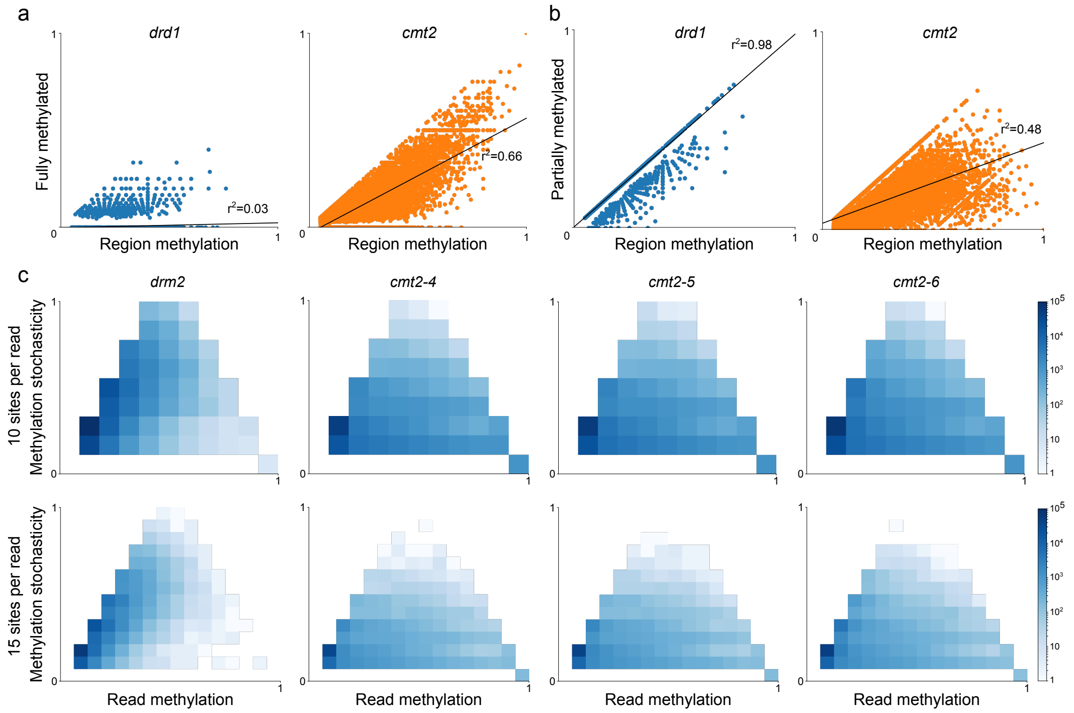


Figure S2: **Meta-analyses of CHH methylation mutant data.** (a-b) Meta-analyses of read methylation data: (a) correlation between fully methylated read proportion of regions and region methylation, (b) correlation between the methylation of partially methylated reads of regions and region methylation. Linear regression shown as black lines with squared correlation coefficient (p-value $< 1 \times 10^{-10}$ for all correlations) (c) Read analysis of *drm2* and *cmt2* mutant samples. Each heatmap plots methylation and stochasticity of reads originating from up to 100 bp regions with 10 (upper panels) and 15 (lower panels) CHH sites.

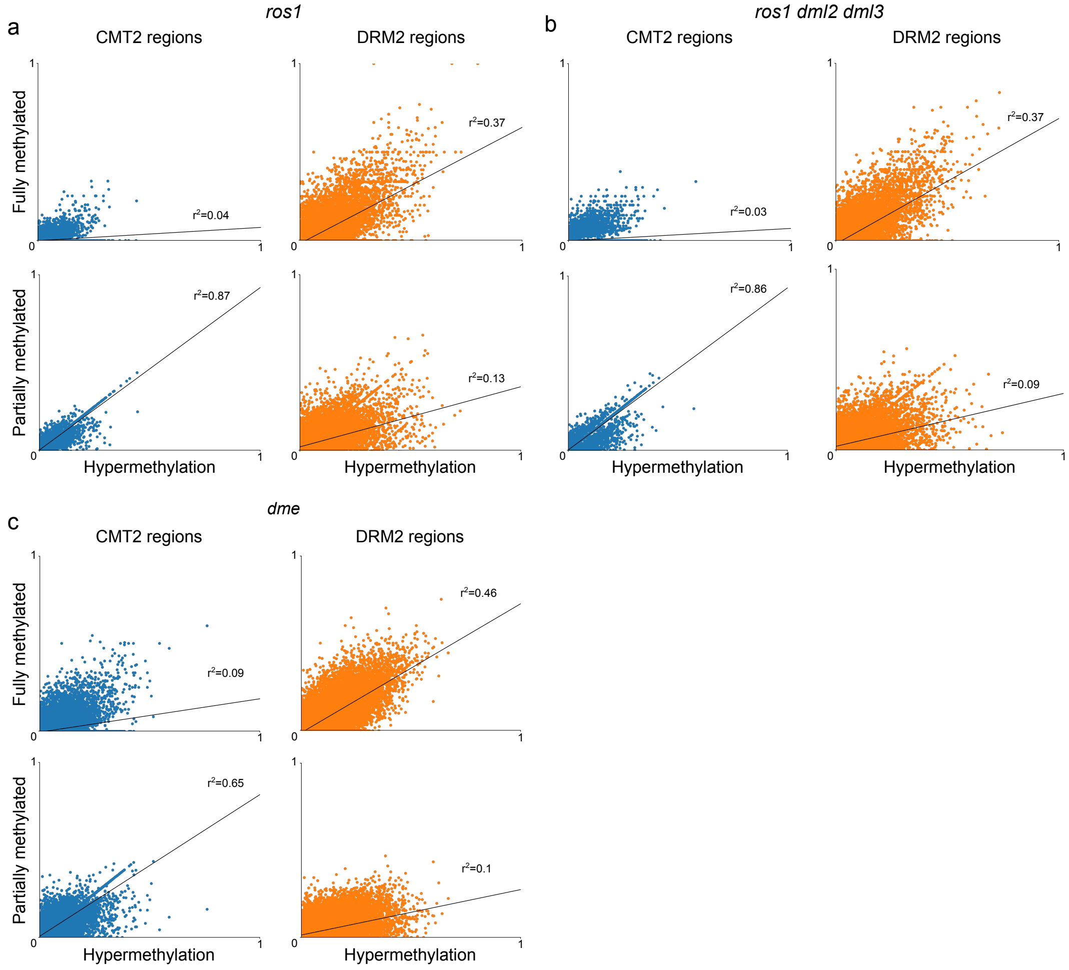


Figure S3: **Meta-analyses of demethylase mutants.** (a-c) Contribution of fully methylated reads (top panels) and partially methylated reads (bottom panels) to the hypermethylation of demethylase mutants *ros1* (a), *rdd* (b) and *dme* (c) compared to their respective wild type samples. Squared correlation coefficients appear next to plotted regression lines (black lines). P-value $< 1 \times 10^{-10}$ for all correlations. In CMT2 regions, demethylase hypermethylation is explained by an increase in partially methylated reads, whereas in DRM2 regions, demethylase hypermethylation is explained by an increase in fully methylated reads.

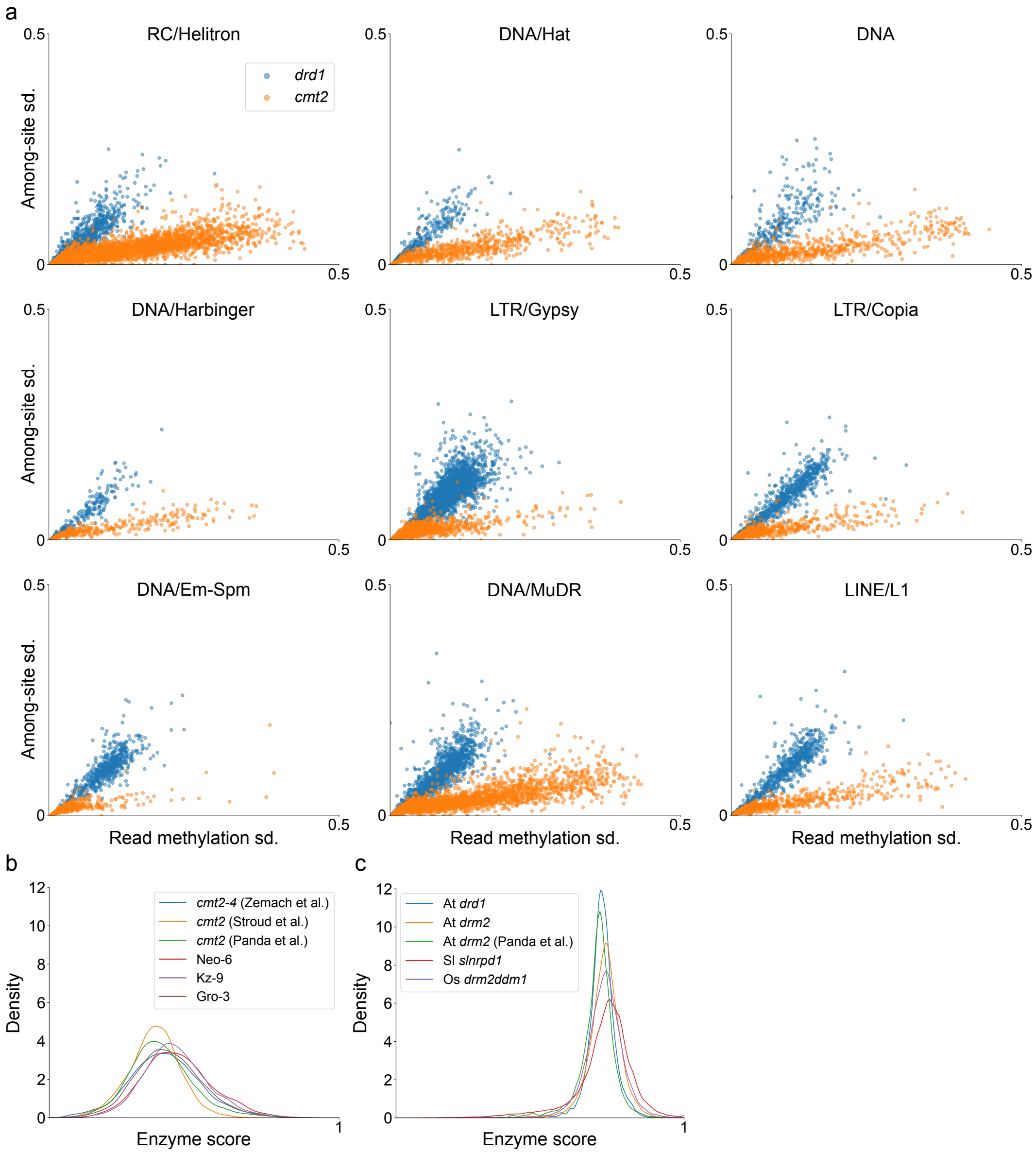


Figure S4: **Analysis of whole transposable elements (TEs) according to methylation variation features and enzyme score in CHH methylation mutants.** (a) Correlation between methylation variation features in TEs in *A. thaliana* CHH methylation mutants. Each point represents a single TE. (b) Enzyme score analysis of additional *A. thaliana cmt2* mutants alongside the *cmt2* mutant used to design the classifier. (c) Distribution of TEs according to enzyme score in DRM2-related mutants from multiple species and tissue types.

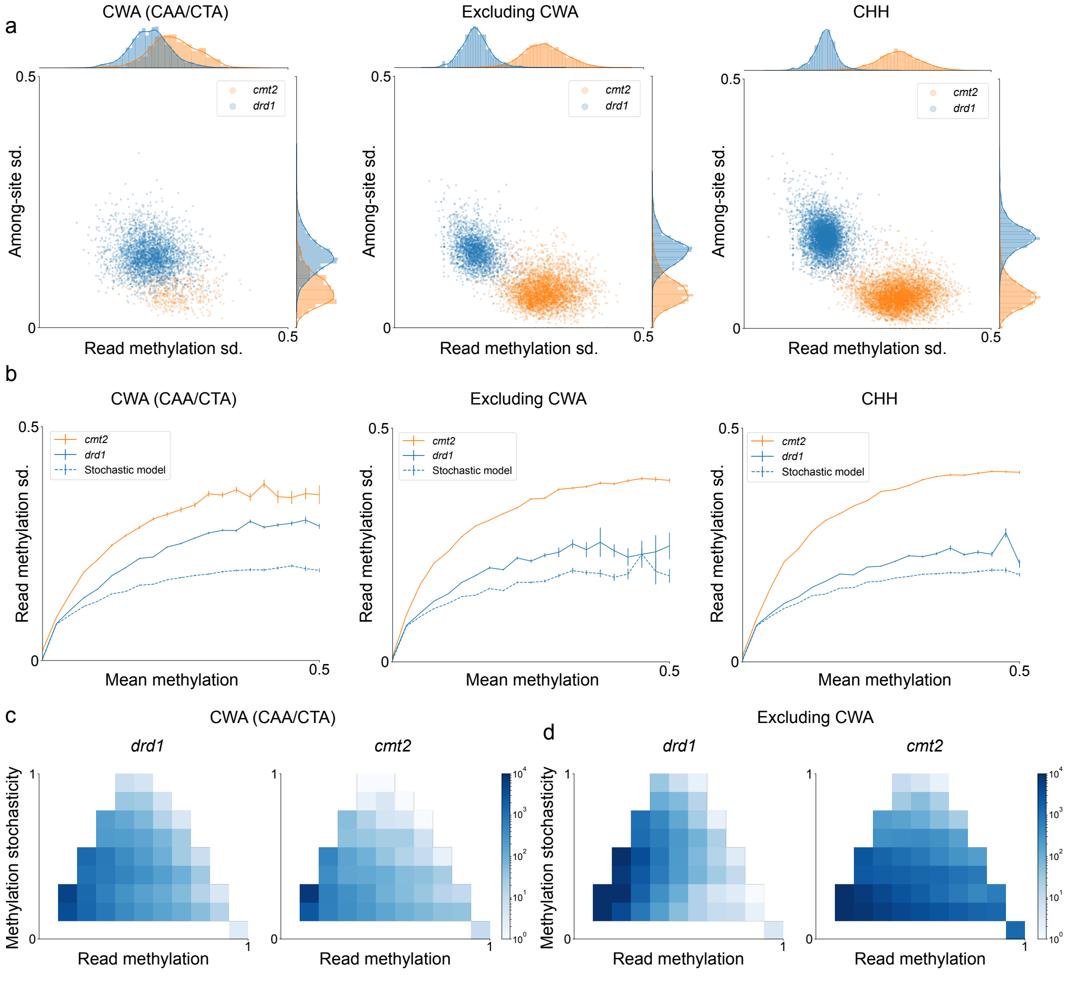


Figure S5: **Sub-context differences in methylation variation.** (a) Separation of TEs by read methylation sd. and among-site sd. in different CHH subcontexts. (b) Relationship between region mean methylation and read methylation sd. in CHH subcontexts in *drd1* and *cmt2*, compared to a stochastic model of read methylation in *drd1* (dashed blue line). Lines represent the average read methylation sd. of a region for a given methylation level. (c-d) Read analysis from CWA-methylated regions (c) and other CHH subcontexts (d) in *drd1* and *cmt2* mutants.

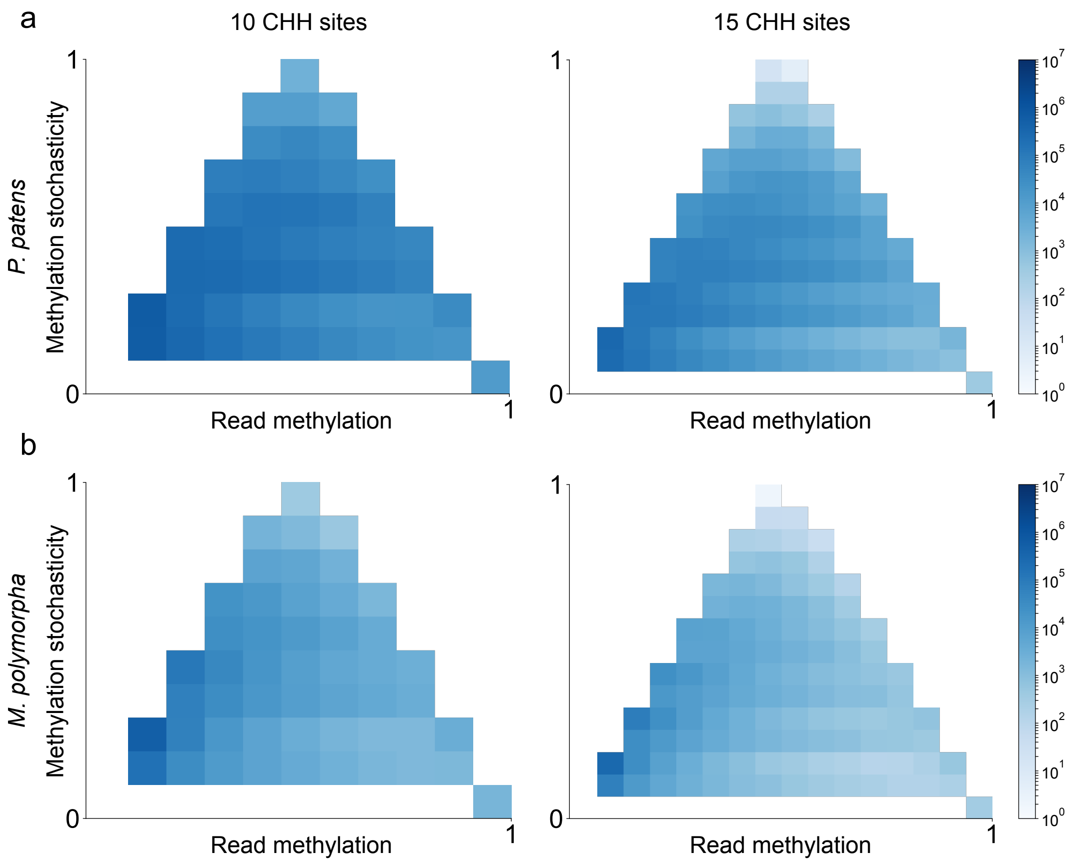


Figure S6: **Read and element-specific patterns of enzyme activity.** (a-b) Read analysis of *P. patens* (a) and *M. polymorpha* (b) which lack a peak in enzyme score aligned to *cmt2* mutants. Each heatmap plots the number of reads in each sample with methylation and stochasticity indicated on the axes.

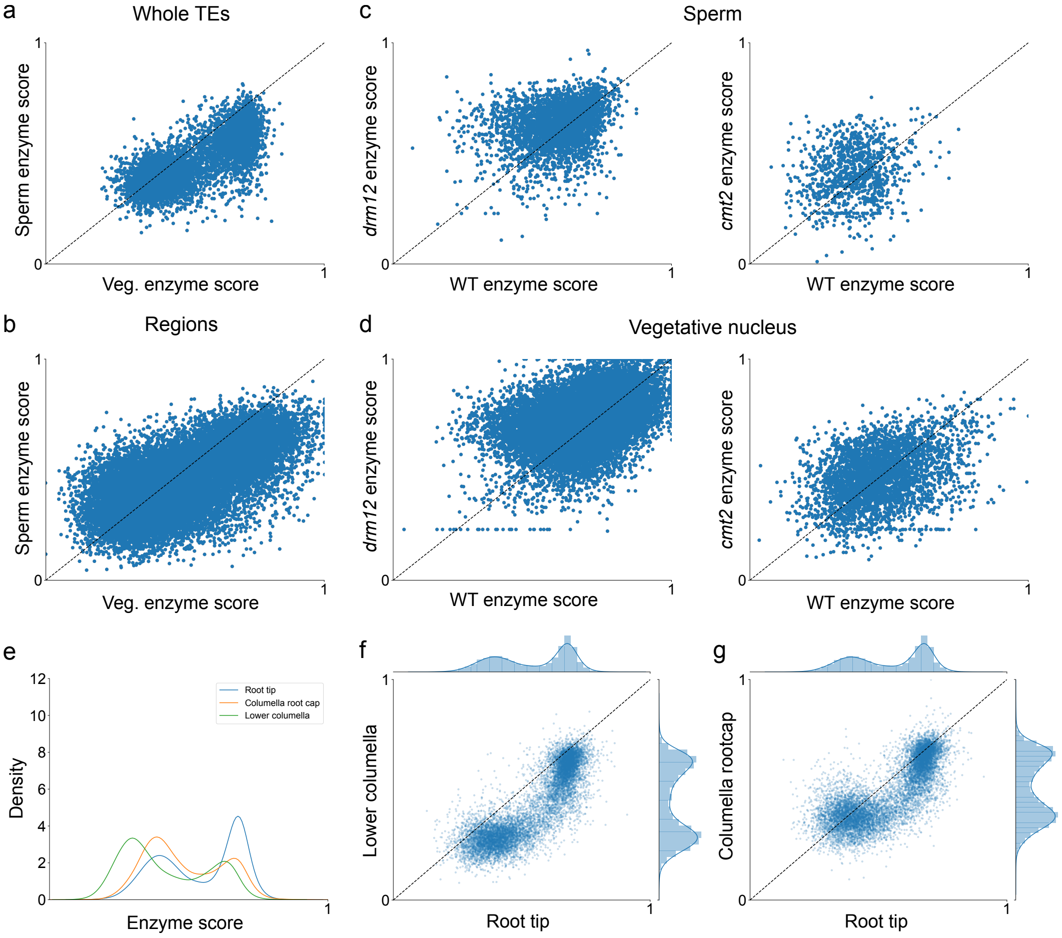


Figure S7: **Tissue-specific patterns of methylation variation.** (a-b) Changes in region enzyme score between *A. thaliana* vegetative nucleus and sperm tissues. Each dot represents an individual TE (a) or CHH-methylated region of up to 30 bp containing 5 CHH sites, plotted according to its enzyme score in each tissue. Only TEs/regions that are methylated ($>= 10\%$) in both samples are plotted. Distance from the dashed line ($f(x) = x$) indicates a predicted change in regulation between tissue types. (c-d) Changes in region enzyme score between mutants of samples presented in (b), by the same region selection parameters. (c) Enzyme score of CHH-methylated regions from sperm *drm12* and *cmt2* mutants compared to wild type. (d) Enzyme score of CHH-methylated regions from vegetative nucleus *drm12* and *cmt2* mutants compared to wild type. (e) Distribution of TEs according to enzyme score in *A. thaliana* root subsamples. (f-g) Changes in TE enzyme score between root tip samples. Each dot represents a single TE, plotted according to its enzyme score in the samples indicated on the axes. Distance from the dashed line ($f(x) = x$) indicates a change in enzyme score between the samples.

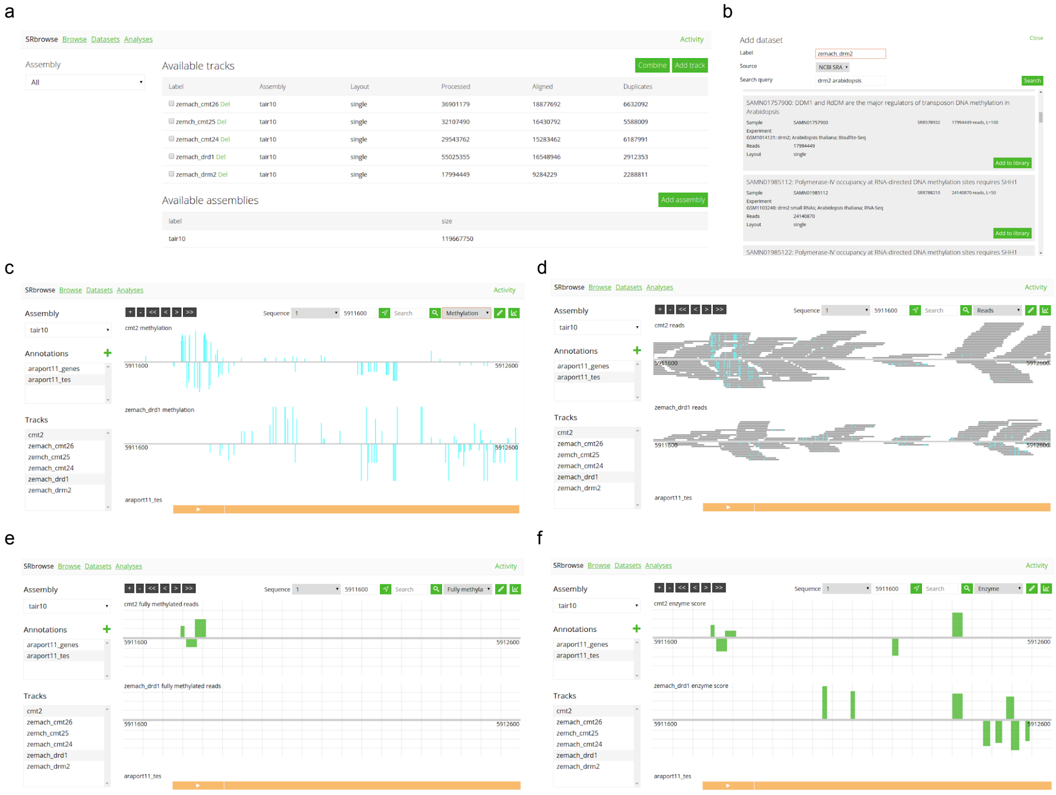


Figure S8: **SRBrowse, a tool for displaying and analysing BS-seq data at single-read resolution.** (a-b) Screens for loading data into the tool, showing currently loaded datasets (a). (b) Adding data into the tool directly from NCBI. (c-f) Genome browser with loaded methylation tracks and TE annotation for *A. thaliana* (the same region is shown in all panels). (c) Methylation signal display of CHH methylation. (d) Display of all reads in a specific sample. Reads are drawn as grey lines, methylation signal within the reads is drawn as cyan boxes. (e-f) Feature signals of regions containing 5 CHH sites with a maximal span of 30bp and minimal methylation of 5%: (e) proportion of fully-methylated reads in region; (f) enzyme score of region.
